# Cerebral perfusion in post-stroke aphasia and its relation to residual language abilities

**DOI:** 10.1101/2022.07.01.498516

**Authors:** Maria V. Ivanova, Ioannis Pappas, Benjamin Inglis, Alexis Pracar, Timothy J. Herron, Juliana V. Baldo, Andrew S. Kayser, Mark D’Esposito, Nina F. Dronkers

## Abstract

Stroke alters blood flow to the brain resulting in damaged tissue and cell death. Moreover, the disruption of cerebral blood flow (perfusion) can be observed in areas surrounding and distal to the lesion. These structurally preserved but sub-optimally perfused regions may also affect recovery. Thus, to better understand aphasia recovery the relationship between cerebral perfusion and language needs to be systematically examined. In the current study, we aimed to evaluate 1) how stroke affects perfusion outside of lesioned areas in chronic aphasia, and 2) how perfusion in specific cortical areas and perilesional tissue relates to language outcomes in aphasia. We analyzed perfusion data from a large sample of participants with chronic aphasia due to left hemisphere stroke (*n*=43) and age-matched healthy controls (*n*=25). We used anatomically-defined regions of interest that covered the frontal, parietal, and temporal areas of the perisylvian cortex in both hemispheres, areas typically known to support language, along with several control regions not implicated in language processing. For the aphasia group we also looked at three regions of interest in the perilesional tissue. We compared perfusion levels between the two groups and investigated the relationship between perfusion levels and language subtest scores while controlling for demographic and lesion variables. First, we observed that perfusion levels outside the lesioned areas were significantly reduced in frontal and parietal regions in the left hemisphere in people with aphasia compared to the control group, while no differences were observed for the right hemisphere regions. Second, we found that perfusion in the left temporal lobe (and most strongly in the posterior part of both superior and middle temporal gyri) and inferior parietal areas (supramarginal gyrus) was significantly related to residual expressive and receptive language abilities. In contrast, perfusion in the frontal regions did not show such a relationship; no relationship with language was also observed for perfusion levels in control areas and all right hemisphere regions. Third, perilesional perfusion was only marginally related to language production abilities. Cumulatively, the current findings demonstrate that blood flow is reduced beyond the lesion site in chronic aphasia and that hypoperfused neural tissue in critical temporoparietal language areas has a negative impact on behavioral outcomes. These results, using perfusion imaging, underscore the critical and general role that left hemisphere posterior temporal regions play in various expressive and receptive language abilities. Overall, the study highlights the importance of exploring perfusion measures in stroke.

## 1. INTRODUCTION

Stroke is a heterogeneous syndrome caused by multiple pathological mechanisms resulting in the disruption of cerebral blood flow (CBF). Decreases in the rate of delivery of blood to brain tissue (measured in units of ml blood/100 g tissue/min) result in subsequent cell death and tissue loss due to the lack of necessary oxygen^1^ leading to motor, cognitive, and language impairments in stroke survivors. In human adults, normal CBF in different gray matter regions typically ranges from 35 to 80 mL/100 g/min depending on various factors such as age, sex, diet, cardiovascular fitness, and health status.^2–5^ CBF is generally considered to be coupled with cerebral glucose metabolism^6,7^, making it an important indicator of tissue functionality. Specifically, in animal models it was shown that brain tissue needs to be perfused at 10% of normal levels to survive, and at least at 30 – 50% for neuronal function (i.e., electrical signaling) to continue.^8,9^ Subsequently, it is expected that lower perfusion may potentially impede normal functioning.^10–12^ Specifically in the context of aphasia, it remains to be answered how CBF after stroke in different brain regions relates to language outcomes and whether it can provide additional insights into the mechanisms of post-stroke recovery.

Research from the acute stroke literature using MRI techniques has demonstrated that CBF is typically disrupted and lowered in perilesional tissue, areas surrounding the neural regions that are permanently damaged.^13–15^ These aberrant perfusion effects persist in the chronic phase, with perilesional perfusion reduced relative to homologous areas in the contralesional hemisphere and similar regions in age-matched healthy controls.^16–20^ The relationship of hypoperfusion of perilesional tissue to language outcomes have been mixed. Some studies have shown that levels of perilesional perfusion are associated with cognitive and language impairments^16,18,21,22^ and recovery of motor function^23^ while others failed to find systematic association between perilesional perfusion and language outcomes.^17,24^

Disruption of perfusion in stroke can also be found beyond perilesional areas. Decreased CBF has been observed in ipsilateral areas distant to the lesion.^5,19,25^ A recent comprehensive study of perfusion in chronic aphasia showed a decrease in perfusion in several areas in the left hemisphere compared to healthy controls (and somewhat unexpectedly an increase in the superior frontal gyrus); however, these differences were not associated with functional language outcomes.^16^ In this study, the left hemisphere ROIs falling within the distribution of the middle cerebral artery were hypoperfused, while regions in the anterior cerebral artery were hyperperfused. Another recent small cohort study employing individualized perfusion cutoffs based on right hemisphere perfusion values demonstrated a strong relationship between hypoperfusion of regions in the left posterior temporal and inferior parietal areas and general language ability and auditory comprehension.^5^ Furthermore, perfusion levels in the middle cerebral artery territory of the left hemisphere have been shown to predict stroke recovery, with initially higher CBF predictive of better treatment outcomes in aphasia.^17,24,26^

A few studies have investigated the effects of perfusion in the right contralateral hemisphere and these have found increased perfusion.^16,17,25^ Thompson and colleagues^16^ proposed that perfusion in the right hemisphere is increased potentially as a form of compensatory autoregulatory process, on the assumption that total blood flow to the brain remains approximately the same post-stroke. It is unclear whether these changes are related to functional outcomes. Thompson and colleagues^16^ found no such relationship. Conversely, Boukrina and colleagues^17^ showed that increased perfusion in the right hemisphere regions that are part of the reading network based on the Neurosynth database^27^ was associated with lower word reading accuracy both in the subacute and the chronic stages (but not related to performance on other language tasks).

The lack of consensus in prior perfusion studies in aphasia may be due to small sample sizes (e.g., typically 1-30 participants) and other methodological issues, such as age differences between the aphasia and the control groups, different imaging sequences and processing algorithms, arbitrary cutoffs, varying definitions of perilesional areas, and different parcellations. Larger group studies using perfusion in chronic post-stroke aphasia are needed to address existing knowledge gaps.

In the current study, we aimed to explore how stroke affects perfusion outside of lesioned areas in chronic post-stroke aphasia, and how perfusion in specific cortical areas and perilesional tissue relate to language outcomes in aphasia. Specifically, we hypothesized that perfusion in stroke patients would be reduced outside of lesioned areas, both in perilesional and distant areas, compared to the right hemisphere and to perfusion in age-matched healthy controls. Also, we hypothesized that perfusion levels specifically in perisylvian areas, regions typically known to support language, but not in other brain areas would correlate with language abilities in post-stroke aphasia. Our current work expands upon previous studies in several important ways. First, we analyze one of the largest samples to date of chronic patients with aphasia scanned with perfusion imaging. Second, we explore regions in both hemispheres as well as systematically investigate perfusion in perilesional areas, examining separately perilesional tissue at varying distances from the lesion. Finally, we explore the contribution of perfusion in multiple regions simultaneously to language abilities while accounting for structural damage.

## 2. METHODS

### 2.1. Participants

Forty-three participants with aphasia (PWA; 29 males, 14 females) following a left hemisphere stroke (M_age_ = 65.5 ± 11.1 years, from 43 to 88 years of age) were recruited for the study. All participants except three were strongly right-handed based on the Edinburgh Handedness Inventory ^28^, with these three participants reporting a right-hand preference but some ambidexterity. All passed screening tests for any hearing and visual deficits and had nativelike proficiency in English prior to their stroke. All participants had suffered a single stroke, except for small (< 2 cm) asymptomatic secondary events, with the most recent incident being at least 3 months prior to testing and scanning (M_time post-onset_ = 52.2 ± 72.3 months). PWA in this sample presented with a wide range of speech and language deficits, some performing within normal limits on the Western Aphasia Battery (WAB^29,30^), but still complaining of residual naming and/or comprehension deficits (see Results section for more information). All patients signed IRB-approved consent forms and were tested in accordance with the Helsinki Declaration.

A group of healthy age-matched controls was also recruited. Twenty-five participants (19 males, 6 females) with no neurological history participated (M_age_ = 61.6 ± 11.3 years, from 41 to 84 years of age). There was no significant difference in age between the control and aphasia groups (*t* (66) = −1.35, *p* = 0.18) or gender distribution (*X^2^* (1, *N* = 68) = 0.56, *p* = .46).

### 2.2. Behavioral assessments

The WAB^29,30^ was administered to evaluate the language abilities of the PWA. Participants were assessed with the ten main language subtests, which contribute to the following subtest scores: Fluency, Information Content, Repetition, Naming, and Auditory Comprehension. Scores from these subtests comprise the WAB Aphasia Quotient (AQ), a general measure of aphasia severity.

### 2.3. MRI data: Acquisition, pre-processing, perfusion data analysis

Structural MRI (T1w) and perfusion data were acquired. The participants were scanned at two different sites with slightly differing protocols described below. To account for possible interscanner variations, we normalized individual perfusion by whole-brain perfusion and used scanning site was as a covariate in all the correlation/regression analyses (see section 2.4 Statistical Analysis).

#### 2.3.1. Data acquisition

##### 2.3.1.1. VA cohort

Veterans Affairs (VA) participants underwent anatomical and ASL scans on a 3T Siemens Verio scanner with a 12-channel phased-array head coil at the VA Hospital in Martinez, CA. For perfusion, a pseudo-continuous arterial spin labeling (pCASL) sequence was used with the following parameters: TR/TE = 4000/12ms, flip angle = 90°, bandwidth = 2.6 KHz/pixel, FOV = 22cm, voxel size = 3.4×3.4×6 mm, slice-selective gradient = 6mT/m, 20 axial slices in ascending sequential acquisition order using echo planar imaging (EPI) readout. The labeling duration was 1470ms with a post labeling delay of 1500ms. 80 images were acquired in the interleaved tag/control order for each subject. Since a separate M0 calibration image was not obtained, per current recommendations^31^, we used the first control image as a calibration image for the analysis of this cohort. One high-resolution anatomical image was acquired for each subject with the scan parameters: MP-RAGE sequence, TR/TE = 2200/1.62ms, TI = 900ms, flip angle = 9°, FOV = 256mm, voxel size 1×1×1 mm, 192 sagittal slices, bandwidth = 340Hz/voxel, GRAPPA factor = 2. Twenty-nine PWA and all twenty-five controls were scanned at this location.

##### 2.3.1.2. UC Berkeley cohort

Berkeley participants underwent anatomical and ASL scans on a Siemens 3T Trio scanner with a 32-channel coil at the Henry H. Wheeler Jr. Brain Imaging Center, UC Berkeley, CA. For perfusion imaging, a pCASL sequence with spiral readout was used with the following parameters: TR/TE = 4600/8.7ms, flip angle = 90°, bandwidth = 400Hz/Pixel, FOV = 25cm, 40 slices, voxel size = 3×3×3 mm, phase encoding gradient = 6mT/m. The labeling duration was 1800ms with a post labeling delay of 2000ms. 16 images were acquired in the interleaved tag/control order for each subject. A stack-of-spirals readout was used with a 4-shot spiral interleave for each of the 3D phase encoding steps with background suppression on.^32^ An equilibrium magnetization (M0) image was also obtained and later used in the kinetic model to compute CBF values. One high-resolution anatomical image was also acquired for each subject with the following parameters: MP-RAGE sequence, TR/TE=2300/2.96ms, TI=900ms, flip angle = 9°, FOV = 256mm, voxel size 1×1×1 mm, 208 sagittal slices, bandwidth = 240Hz/voxel, GRAPPA factor = 2. Fourteen PWA were scanned at this location.

#### 2.3.2. ASL data preprocessing

The processing steps were identical for the two cohorts. Tag-control images were motion corrected using FSL’s function MCFLIRT ^33^. Cerebral blood flow (CBF) maps were obtained using FSL’s command ‘oxford_asl’ on the motion corrected ASL data^34^ with the parameters tailored to reflect each site’s acquisition parameters. CBF maps were quantified in standard physiological units (ml blood/100mg tissue/min) using a standard kinetic model.^31^ Labeling efficiency was set to α=0.72 and the longitudinal relaxation time of the blood was set to T1_b=1650ms. No further smoothing was performed.

#### 2.3.3. Lesion segmentation and structural data preprocessing

The participants’ lesions were traced directly onto the patient’s native T1-weighted images using MRIcron software^35^ by trained research assistants, and then reviewed and verified by MI, IP, and ND. Next, we used the ANTs toolbox^36^ to perform brain extraction on the structural T1s. We also used ANTs to segment the T1 images and obtain probability maps corresponding to different tissues (gray matter, white matter, and cerebrospinal fluid (CSF)). We then performed the following registrations to be able to bring regions of interest (ROIs) to native CBF space. (*A*) The CBF maps were registered to the brain extracted images (T1 native space) using an affine transformation. (*B*) The brain-extracted T1 images were registered to the MNI-152 space. To do so we used ANTs with a transformation that consists of an initial rigid plus affine transformation followed by a diffeomorphic, “SyN” transformation, while cost-function masking the lesion. We used the MNI152NLin2009cAsym version of the MNI image as the target MNI image.^37^ Using the inverse of the transformation obtained in (A), we were able to map individually-defined lesion masks, as well as perilesional ROIs, from native T1 space to CBF space. Using the combined inverse transformations of (*A*) and (*B*) we were able to map atlas-based ROIs defined in MNI-152 to native T1 and then to CBF space. In the analysis we use the following ROIs:

i. *Atlas-based ROIs* – were taken from the Harvard-Oxford Cortical Structural Atlas in FSL.^33,38^ The segmentation from this atlas was selected as it contains ROIs of optimal size. Given the low-resolution of the perfusion data we needed ROIs large enough to minimize partial volume effects, but small enough to ensure adequate spatial specificity for distinct language regions. From the atlas we selected 11 ROIs that covered Perisylvian language regions (Inferior frontal gyrus (IFG) triangularis, IFG opercularis, Supramarginal anterior, Supramarginal posterior, Angular gyrus, Temporal pole, Superior temporal gyrus (STG) anterior, STG posterior, Middle temporal gyrus (MTG) anterior, MTG posterior, MTG temporal-occipital). We were interested whether perfusion levels in those ROIs would be related to residual language abilities. We also included four control ROIs (Frontal pole, Central, Superior frontal gyrus (SFG), Occipital pole) that covered regions typically not related to language processing. See Figure 1 for representation of these ROIs. These ROIs in MNI-152 space were brought to native CBF space using the combination of transformations described previously in (*A*) and (*B*).
ii. *Perilesional ROIs* – were obtained by eroding the lesion mask in native T1 space to 5mm and subtracting it from the original lesion mask. This process was repeated stepwise to obtain additional perilesional masks for 5-10mm and 10-15mm bands outside the lesion. These three perilesional ROIs were brought to CBF space using the inverse affine transformation described previously in (*A*).

**Figure 1.**
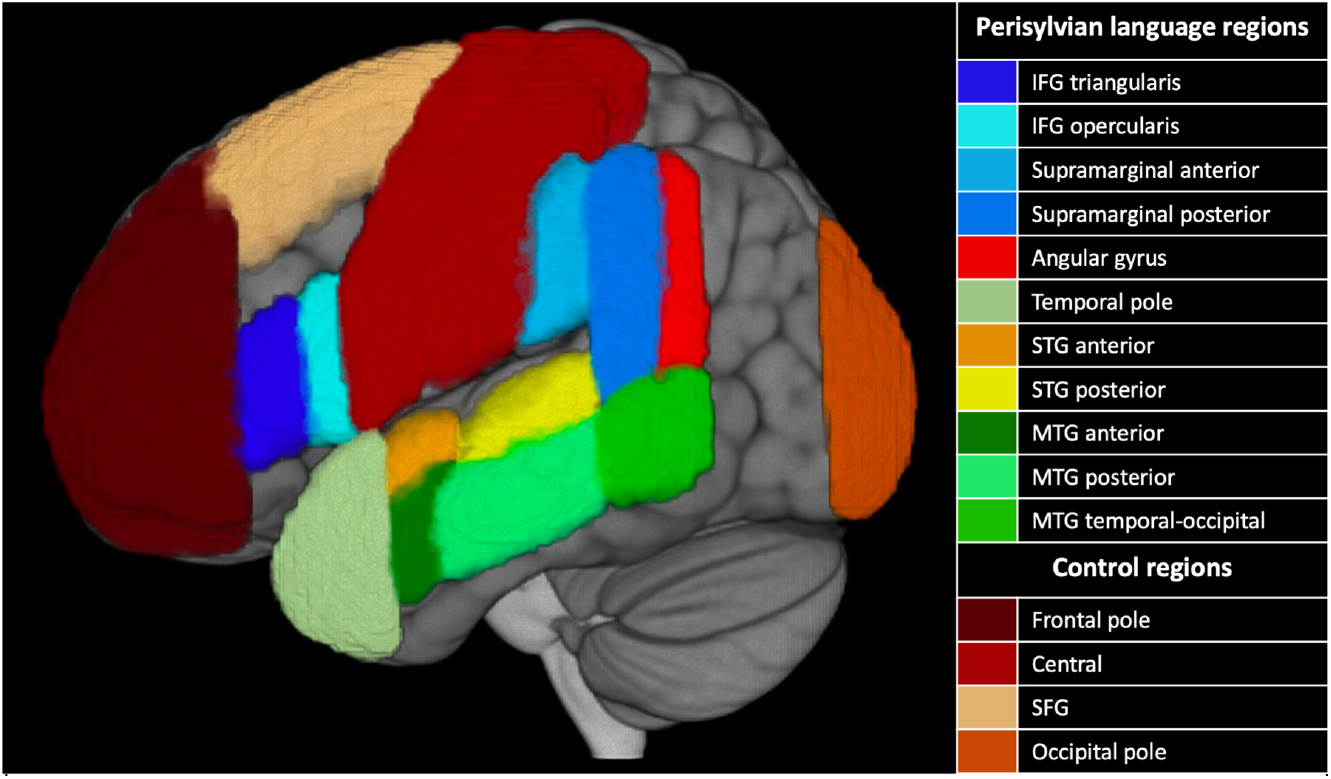
Representation of Atlas-based ROIs taken from the Harvard-Oxford atlas used in the current analysis. IFG – inferior frontal gyrus, STG – superior temporal gyrus, MTG – middle temporal gyrus, SFG – superior frontal gyrus.

#### 2.3.4. Region of interest (ROI) analysis of ASL data

For all the ROIs mean CBF values were obtained as the average CBF within each ROI. Because some ROIs overlapped with regions of no interest (e.g., perilesional ROIs could end up covering the ventricles or include voxels outside the brain), we required the CBF values of each ROI to exclude any CSF values or values outside the brain mask for all our calculations. Right hemisphere areas were also excluded from all perilesional ROIs. In addition, for the PWA cohort, lesioned voxels determined by the lesion mask were excluded from all the atlas-based ROIs at the participant level. Finally, lesion load was defined as the percentage of voxels in the ROI covered by the lesion.

### 2.4. Statistical analysis

For all the statistical analyses, we divided perfusion values by the whole-brain perfusion signal (mean perfusion across the whole brain with the lesion site masked). This approach allowed to account for individual variability in tagging efficiency and other potential session-specific and scanner-specific artifacts. To control for multiple comparisons, we adjusted the *p*-value in each analysis by the number of atlas-based ROIs in each hemisphere. Thus, our critical significance threshold was *p* < .0033 (0.05/15; two-tailed) for both between-group comparisons and correlation analyses. All data analyses were performed in R ver. 4.1.2.^39^ and figures were drawn in ggplot2 ver. 3.3.5.^40^

For between-group comparisons of perfusion levels in different ROIs, the following procedure was implemented. First, we checked whether perfusion data for a specific ROI for each group followed a normal distribution (Shapiro test). If the data were normally distributed in both groups for that ROI, we ran an F-test to test for homogeneity of variances. For those ROIs where all these assumptions were satisfied, we ran the standard independent samples t-tests to compare perfusion levels between groups. If the assumption of homogeneity of variances was violated, an unequal variances t-test (Welch’s test) was performed. If in both or in one of the groups the data were not normally distributed, then the non-parametric two-samples Wilcoxon rank test was performed. This procedure was implemented for comparing PWA to age-matched controls and for checking whether perfusion levels differed between those PWA with a lesion in a given atlas-based ROI and those without that lesion. Also, to explore the relationship between perfusion levels and lesion status, we performed correlations between the two metrics – mean adjusted perfusion and lesion load – for each ROI.

For within-group comparisons of adjusted perfusion levels in left hemisphere ROIs to homologous right hemisphere ROIs, we first verified that the differences between homologous left and right ROIs were normally distributed. If that assumption was satisfied, then a standard paired samples t-test was run. If not, then we performed a paired two-samples Wilcoxon test.

Finally, to analyze the relationship between language measures and perfusion levels, we first performed a partial correlation analysis between mean adjusted perfusion in an ROI and language metrics, accounting for age, gender, time post-onset, scanning site, and lesion volume. In addition to using the omnibus lesion volume, we accounted for lesion load to that specific ROI as well. Partial correlation analysis was done with the package ppcor.^41^ This analysis was followed by a regularized lasso regression, so that we could (1) analyze the impact of perfusion in all the ROIs simultaneously, while controlling for lesion load to the respective ROIs; and (2) outline in which regions perfusion levels were associated with different language abilities. Regularized regression allows determination of salient relationships in complex datasets and has been successfully used previously in neuroimaging studies to select relevant neural predictors.^42^

Lasso regression (alpha = 1) differs from traditional multiple linear regression as it employs L1-normalization to regularize model coefficients so that unimportant features are eliminated, preventing overfitting and improving generalization on test data.^43,44^ This approach is also recommended for instances when the number of cases is comparable to the number of predictors, as in the present case. The regularization term provides a constraint on the size of weights, and it is controlled by parameter λ, with larger λ leading to more shrinkage. When there are many correlated variables in an ordinary linear regression model, the weights can be poorly determined and exhibit high variance. Using λ to impose a size constraint on the coefficients alleviates this problem, and a lasso (L1) regularized regression specifically will assign beta weights of 0 to weak predictors.^43^ In this analysis, we included all the covariates from the partial correlation above and all the perilesional and atlas-based ROIs. Predictors first had to be standardized, so that their absolute values would not influence the weights. Next, an optimal λ was selected through leave-one-out cross-validation, with the λ that minimized residual mean squared error in the model used in the analysis. These regression analyses were performed with the glmnet package.^45^

### 2.5. Data availability

The datasets presented in this article are not readily available: Research data are not shared per Department of Veteran Affairs privacy regulations. Requests to access the datasets should be directed to ND.

## 3. RESULTS

### 3.1. Differences in perfusion levels between groups

Lesion overlays for the aphasia group are presented in Figure 2. Maximal lesion overlap was observed in frontal and subcortical temporoparietal areas.

**Figure 2.**
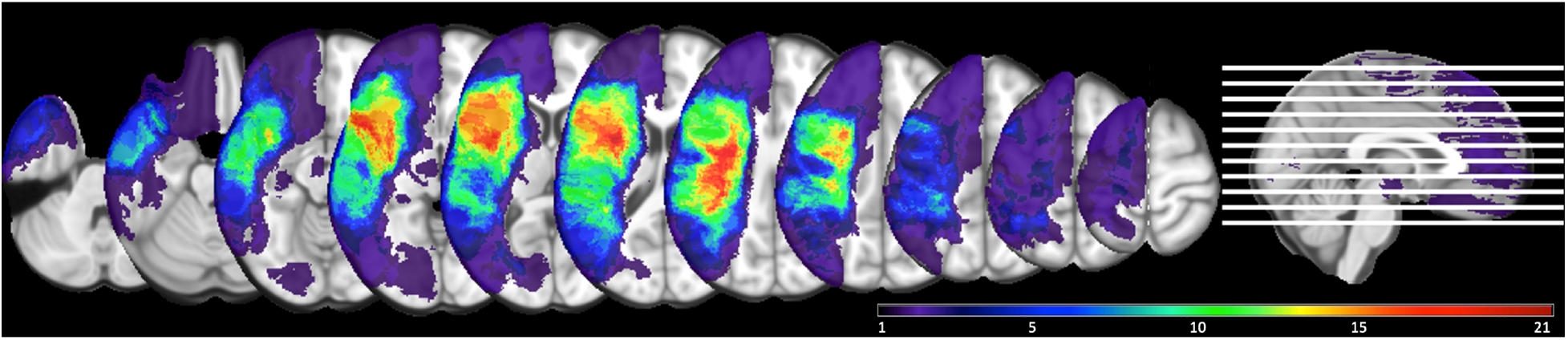
Lesion overlay map (*n =* 43) demonstrating overlap across participants’ lesions, with brighter colors indicating greater number of patients having a lesion in each voxel (ranging from a minimum of one participant’s lesion in a voxel and a maximum of 21).

Raw and adjusted (by the whole-brain perfusion) perfusion values for left and right hemisphere ROIs for the aphasia and the control groups are presented in Figure 3A and 3B respectively (see Appendix, Tables A1 and A2 for actual values and test statistics). For this and all the subsequent analyses, lesioned voxels were excluded from the calculation of perfusion in the left hemisphere. As can be seen from Figure 3A, the majority of the left hemisphere ROIs and some of the right hemisphere ROIs showed significantly lower raw perfusion values in PWA relative to controls as determined by independent samples t-tests/Wilcoxon rank tests. Wholebrain perfusion was also significantly lower in PWA compared to controls (*M*_Controls_ = 32.4, *M*_PWA_ = 26.2; *t* (66) = 3.42, *p* = .001). For the adjusted perfusion values (Figure 3, panel B), only left ROIs, specifically regions in the frontal and parietal regions, showed significantly decreased perfusion in the PWA group compared to age-matched controls. No differences were observed in the right hemisphere. Comparison of raw and adjusted perfusion values for the participants just from the VA cohort yielded largely similar findings, indicating between group differences in adjusted perfusion for the left frontal and parietal regions (see Appendix, Figure A1 for details). For all subsequent analyses adjusted perfusion values are used.

**Figure 3.**
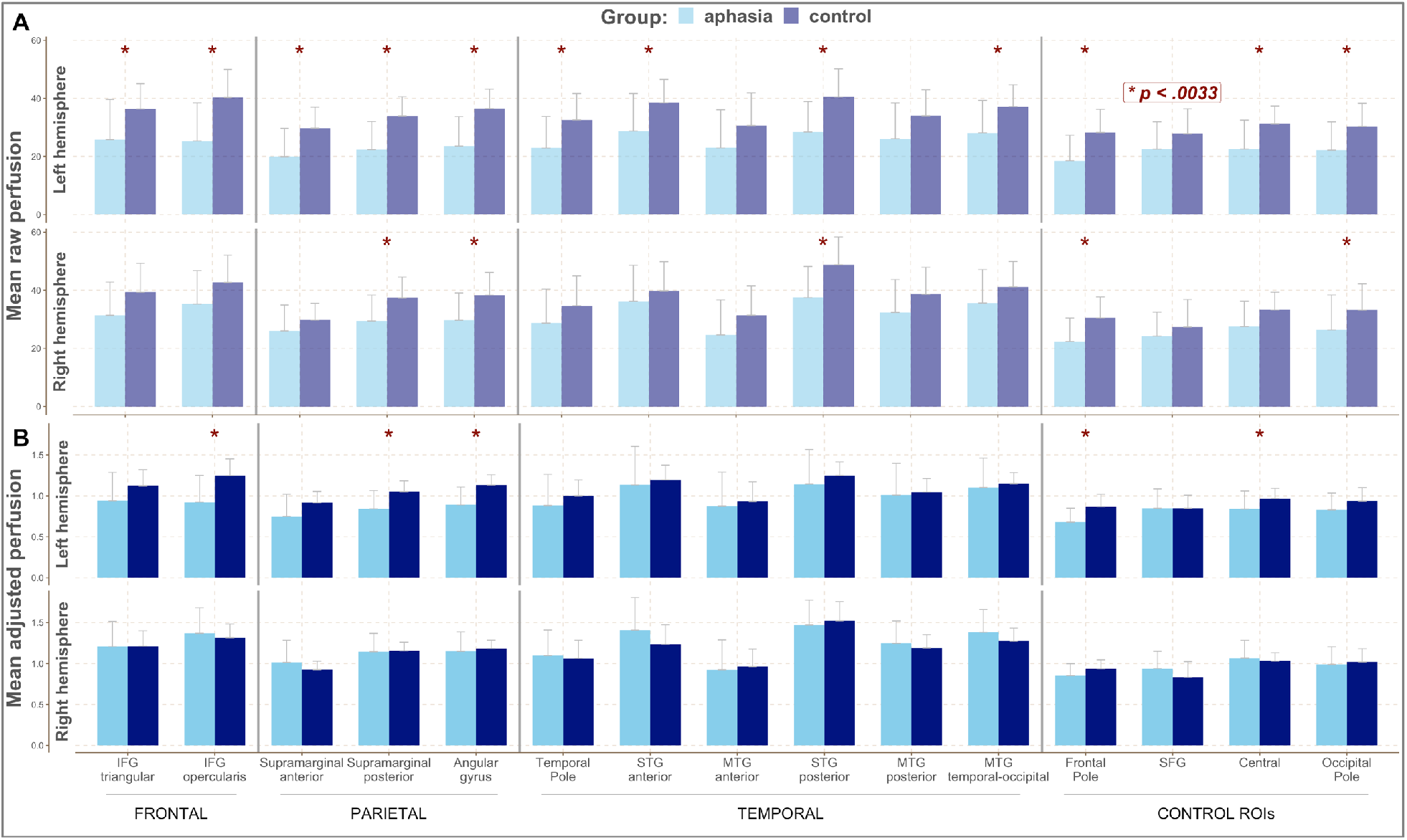
Mean perfusion values for the aphasia and the control groups across left and right hemisphere ROIs. Red asterisks mark significant differences between groups for a given ROI. *Panel A* – Mean raw perfusion. *Panel B* – Mean adjusted perfusion. IFG – inferior frontal gyrus, STG – superior temporal gyrus, MTG – middle temporal gyrus, SFG – superior frontal gyrus.

### 3.2. Differences within groups: comparing perfusion levels in homologous regions in the left and right hemispheres

Next, we compared perfusion levels in the left hemisphere ROIs to homologous regions in the right hemisphere (see Figure 4, panel A) for both the control and aphasia groups. There was a statistically significant decrease in perfusion across all regions of the left hemisphere in the aphasia group, except the anterior part of the MTG and SFG. The control group showed interhemispheric differences for a limited set of regions, including the posterior portion of the supramarginal gyrus, the posterior part of the temporal lobe, and the area around the central sulcus, with the left ROIs having lower perfusion compared to their right hemisphere counterparts. Additionally, we compared the extent of this asymmetry (differences between perfusion in the right vs. left hemispheres) between the two groups (see Figure 4, panel B). Asymmetry of perfusion was more pronounced in the PWA group compared to the control group for frontal and parietal regions, indicating that the PWA group showed greater differences between perfusion in homologous ROIs specifically in these areas.

**Figure 4.**
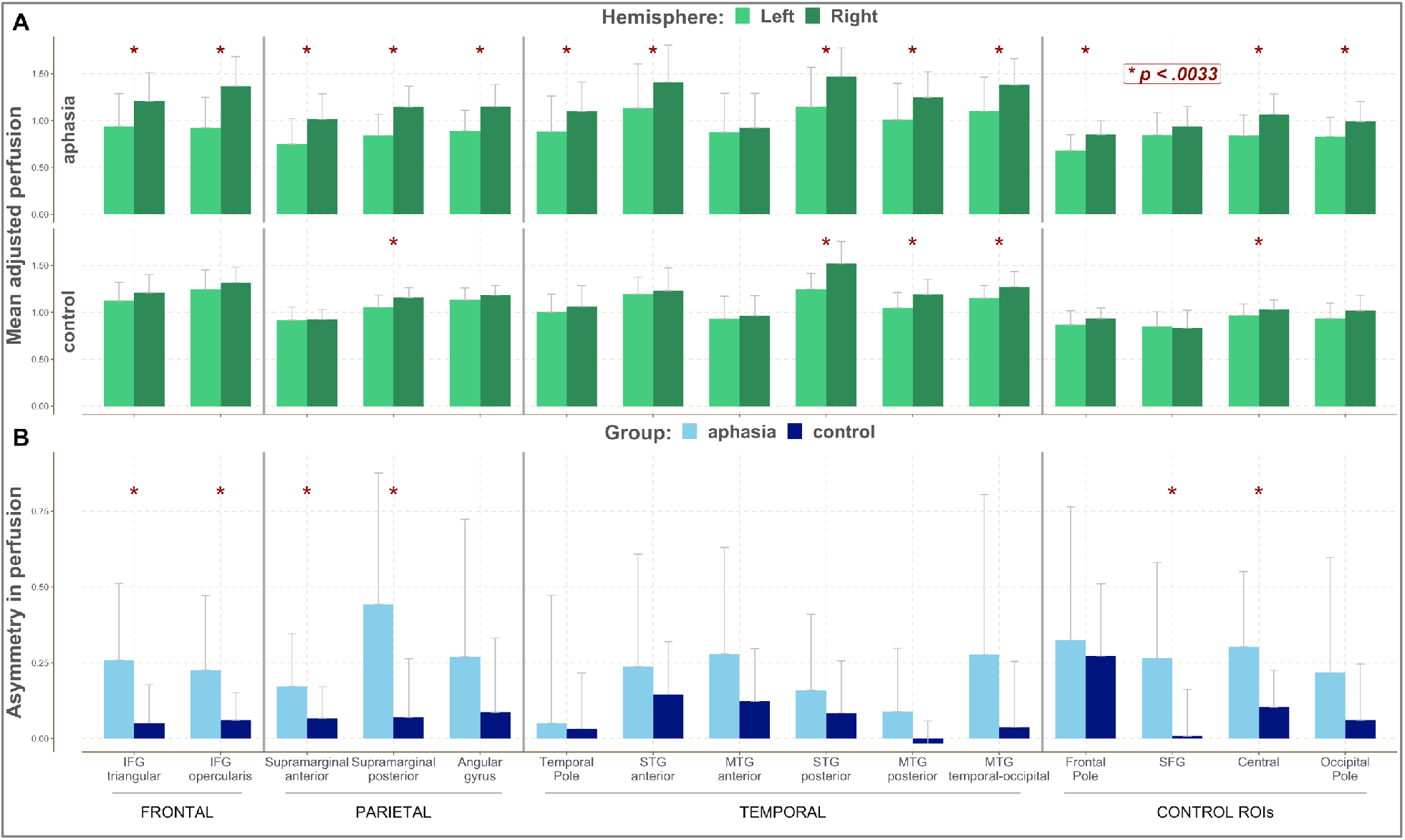
Comparison of perfusion in the left and right hemisphere ROIs between the two groups. Red asterisks mark significant differences. *Panel A* – Mean adjusted perfusion for the left and right hemisphere ROIs in the aphasia and the control groups. *Panel B* – Asymmetry of right-left hemisphere perfusion between the aphasia and the controls groups. IFG – inferior frontal gyrus, STG – superior temporal gyrus, MTG – middle temporal gyrus, SFG – superior frontal gyrus.

### 3.3. Impact of lesion on perfusion levels

Next, we investigated whether perfusion levels were impacted by having a lesion in a specific ROI. For each ROI, we split participants into two groups – those in whom this ROI was lesioned in the left hemisphere (lesion load greater than 0) and those in whom it was spared (lesion load equal to 0). We then compared the perfusion levels between these two groups in that specific ROI in both hemispheres. This procedure was repeated across all the ROIs. In other words, for each comparison we regrouped the participants based on their lesion load for that specific ROI in the left hemisphere. Quantitatively perfusion was higher in the ‘Spared ROI’ group compared to the ‘Lesioned ROI’ group across the left perisylvian regions, but the difference was statistically significant only for the IFG opercularis and different parts of the MTG (see Figure 5). When the same analysis was performed for homologous right hemisphere ROIs no significant or systematic differences were observed between the two groups. In other words, a lesion in the left hemisphere had no impact on perfusion in the homologous right hemisphere regions.

**Figure 5.**
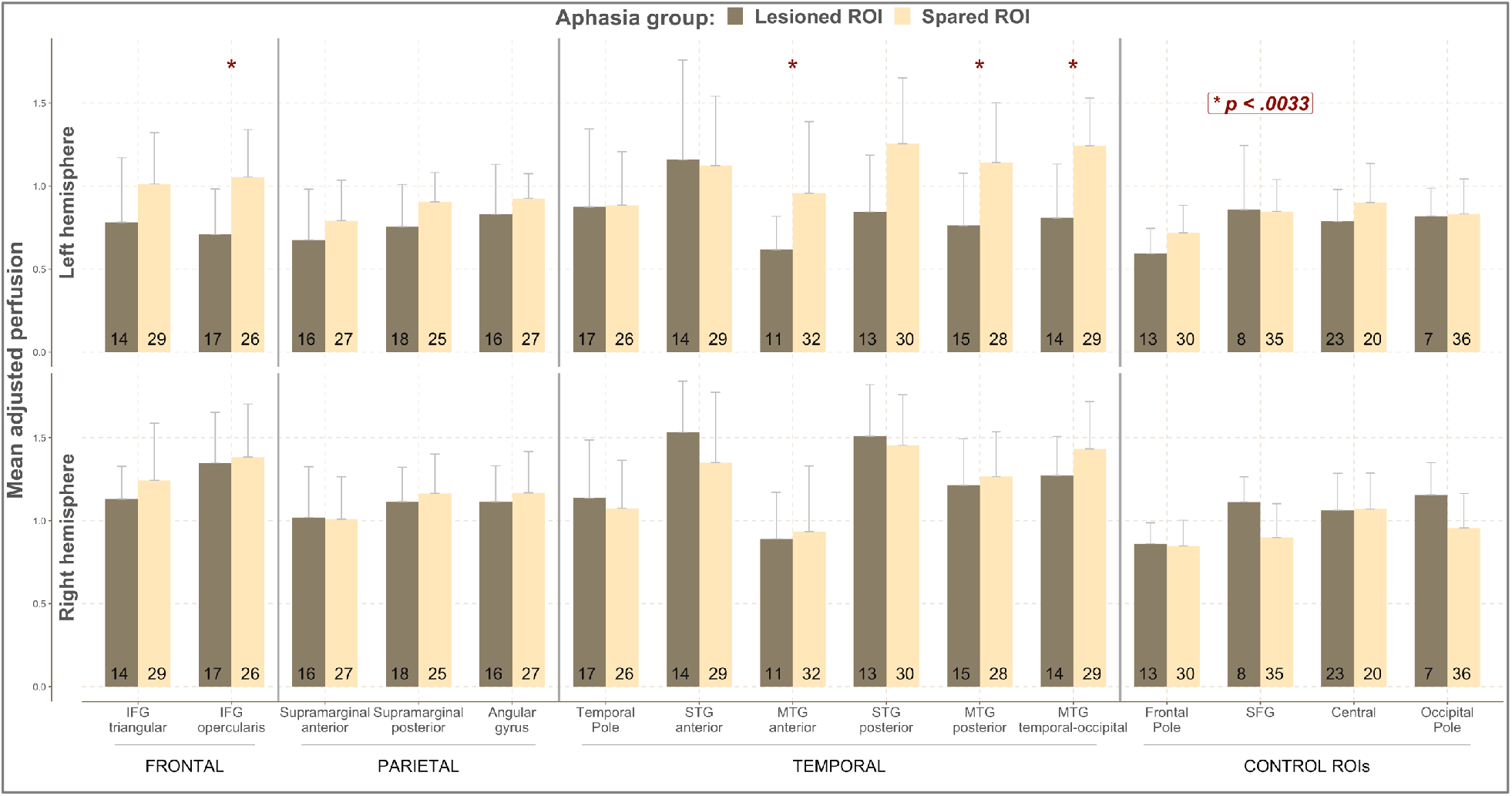
Mean adjusted perfusion for the left and right hemisphere ROIs in the aphasia group with (‘Lesioned ROI’) and without a lesion (‘Spared ROI’) in a given ROI in the left hemisphere. For each ROI for both hemispheres participants are grouped into two subgroups – those in whom this ROI was lesioned in the left hemisphere and those in whom it was spared. The number in each column indicates the number of participants in each subgroup. Red asterisks mark significant differences between the two aphasia subgroups. IFG – inferior frontal gyrus, STG – superior temporal gyrus, MTG – middle temporal gyrus, SFG – superior frontal gyrus.

Further, we explored the relationship between perfusion levels and lesion location by correlating perfusion levels across the different left hemisphere ROIs with their respective lesion load. Non-parametric spearman correlations were performed as the lesion load data was not normally distributed. Significant correlations were only detected for the IFG opercularis (*r* = −.57, *p* < .001), posterior (*r* = −.48, *p* = .001) and temporal-occipital part of the MTG (*r* = −.6, *p* < .001). This indicated that, in these left hemisphere ROIs, having a higher lesion load was associated with having lower perfusion levels in the spared parts of these regions.

Finally, we compared perfusion in the three different perilesional ROIs. Perfusion in the perilesional 0-5mm band (*M*_0-5mm_=0.86 ± 0.19) was significantly lower than in the 5-10mm band (*M*_5-10mm_=0.96 ± 0.16; *t* (42) = −7.5, *p* < .001), which in turn was lower than in the 10-15mm band (*M*_10-15mm_=1.01 ± 0.12; *t* (42) = −3.14, *p* = .003). Only perfusion in 0-5 mm band was significantly lower compared to whole-brain perfusion (*t* (42) = −4.9, *p* < .001).

### 3.4. Correlation between language measures and perfusion levels

Mean WAB subtest scores for the PWA group are presented in Table 1. Note that while some participants had WAB AQ scores above the 93.8 cut-off for aphasia, they continued to experience minor language deficits, particularly for word finding.

**Table 1.**
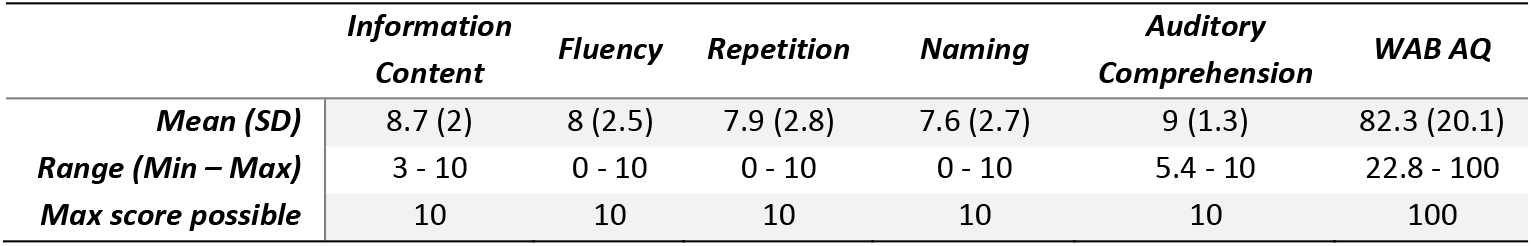
Descriptive statistics for WAB subtest scores.

Partial correlations were performed between language measures and perfusion levels in different atlas-based and perilesional ROIs in the aphasia group accounting for relevant demographic (age, sex, time post-onset, scanning site) and lesion variables (lesion volume or lesion load to a particular ROI). In Panel A of Figure 6, correlations between language and perfusion metrics accounting for demographic variables and lesion volume are presented, while in Panel B, correlations between language and perfusion metrics accounting also for lesion load to individual ROIs are shown. Note that the latter analysis is not applicable to perilesional ROIs, as perilesional bands by definition are not lesioned in any participants. No significant correlations were observed between perfusion levels in the right hemisphere ROIs and language abilities even prior to corrections for multiple comparisons. Accordingly, Figure 6 only includes left hemisphere regions.

**Figure 6.**
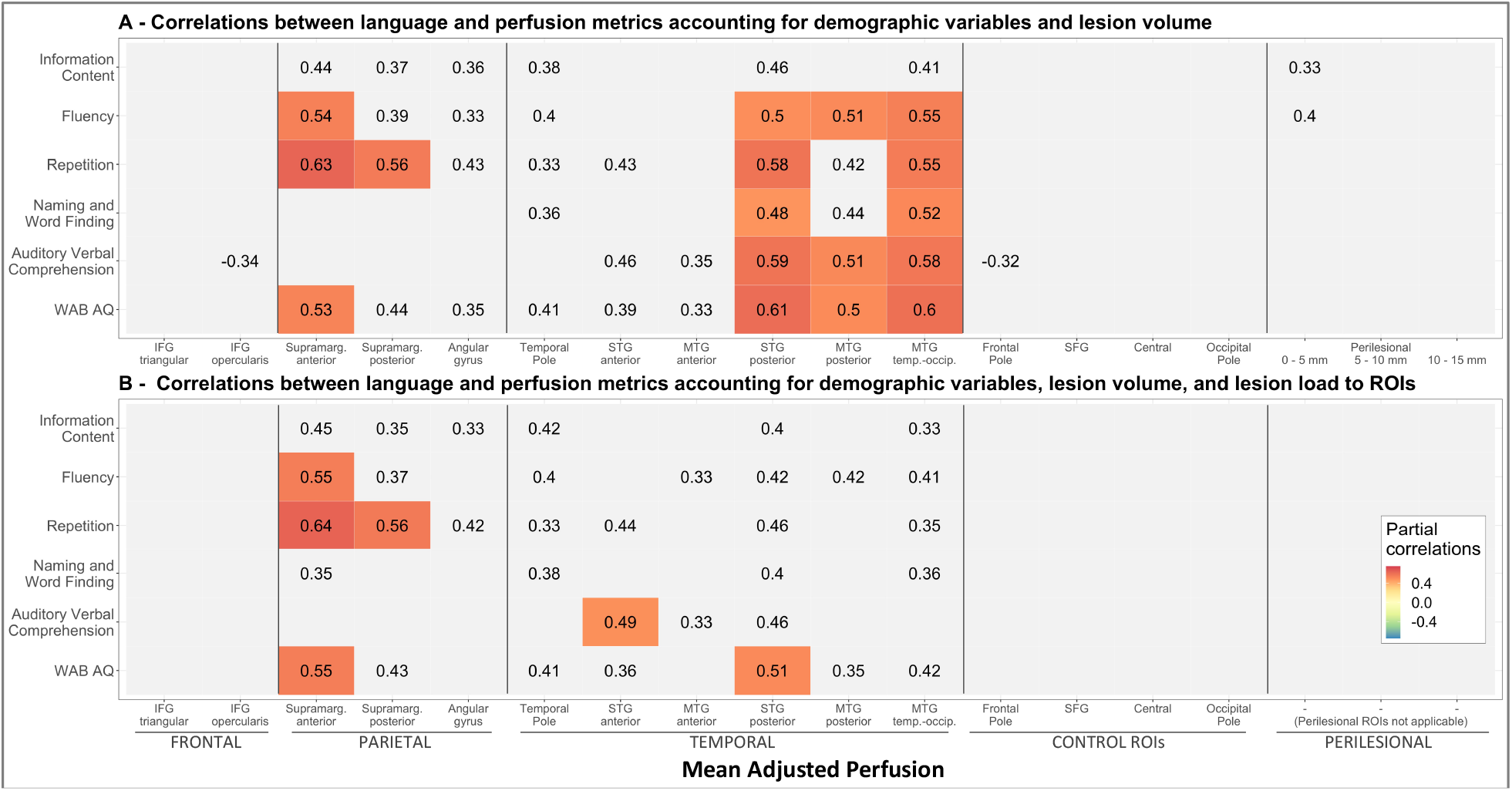
*Panel A:* Partial correlations between language and perfusion metrics in left hemisphere ROIs accounting for age, gender, time post-onset, scanning site, and lesion volume. *Panel B:* Partial correlations between language and perfusion metrics in left hemisphere ROIs accounting for age, gender, time post-onset, scanning site, lesion volume, and lesion load to individual ROIs. Correlations significant at *p* < .05 are printed, while significant correlations after adjusting for multiple comparisons (*p* < .0033) are color coded according to their values. IFG – inferior frontal gyrus, STG – superior temporal gyrus, MTG – middle temporal gyrus, SFG – superior frontal gyrus.

As can be seen from Figure 6 Panel A, correlations remained significant and passed the threshold for multiple comparisons for inferior parietal areas (supramarginal gyrus) and posterior temporal areas (both STG and MTG) when only lesion volume was accounted for. The analysis accounting for lesion load (Panel B) after correction for multiple comparisons did not yield the same pattern of significant correlations between language measures and perfusion levels in the posterior temporal regions. The strength of the association was likely diminished as specifically in those posterior temporal areas there was a significant relationship between lesion load and perfusion levels, as shown previously. Subsequently when lesion load was accounted for (Panel B) only correlations with the supramarginal gyrus remained significant, and limited correlations between STG and language comprehension and general aphasia severity. Further, perfusion levels in perilesional areas were only marginally related to expressive language abilities and did not survive the correction for multiple comparisons. Overall, the data show that perfusion levels in areas beyond the lesion site in the left hemisphere significantly impacted performance on subtests of the WAB even after accounting for demographic and lesion variables (lesion volume and lesion load), while perfusion in the right hemisphere was not associated with residual language abilities.

### 3.5. Lasso regression

While the correlation analysis above provides important insights into the individual role of perfusion in different ROIs to residual language abilities, it does not demonstrate which regions contribute jointly to the observed outcomes and does not outline which regions are most salient for language. To investigate the joint impact of both perfusion and lesion load in different ROIs in the left hemisphere on language outcomes simultaneously, a regularized lasso regression was performed with each of the WAB subtest scores as the outcome variable. Respective regression coefficients are presented in Table 2, with higher values indicative of a stronger relationship with language measures and zero representing lack of such a relationship. Overall, the predictors included in the model accounted for a substantial amount of variance (*R^2^* ranged from .5 to .8), except in the case of repetition where the model accounted for only 17% of the variance and only identified contributions of parietal areas. For the other language abilities, the analyses identified several ROIs in the parietal and temporal lobes simultaneously contributing to language processing, with the supramarginal gyrus and the posterior temporal areas being the strongest predictors. The supramarginal gyrus was particularly strongly related to fluency, while the posterior MTG to naming and auditory comprehension. Only perilesional tissue in the 0 – 5mm band showed a small relationship with expressive language abilities. For the control regions, only the SFG was marginally associated with Fluency. As expected, perfusion in other control areas showed no relationship with language scores.

**Table 2.** Results of Lasso regression analysis with WAB subtest scores as the dependent variables and perfusion in left hemisphere ROIs along with demographic variables and lesion load in left hemisphere ROIs as predictors.

|  |  | **Information**<br>**Content** | **Fluency** | **Repetition** | **Naming** | **Auditory**<br>**Comprehension** | **WAB**<br>**AQ** |
| --- | --- | --- | --- | --- | --- | --- | --- |
| **Covariates** | Lesion volume | -0.176 | 0 | -0.169 | 0 | -0.081 | -0.176 |
|  | Age | 0 | 0 | -0.052 | 0.082 | 0 | 0 |
|  | Time post onset | 0.143 | 0 | 0.153 | 0.256 | 0 | 0.143 |
|  | Scanning site | 0 | 0 | 0 | 0.058 | 0 | 0 |
|  | Gender | 0 | 0 | 0 | 0.061 | 0 | 0 |
| **Mean adjusted perfusion** | IFG triangular | 0 | 0 | 0 | 0 | 0 | 0 |
|  | IFG opercularis | 0 | 0 | 0 | 0 | 0 | 0 |
|  | Supramarginal anterior | 0.222 | 0.051 | 0 | 0.033 | 0.129 | 0.222 |
|  | Supramarginal posterior | 0 | 0.048 | 0 | 0 | 0 | 0 |
|  | Angular gyrus | 0 | 0 | 0 | 0 | 0 | 0 |
|  | Temporal pole | 0.053 | 0 | 0.021 | 0 | 0 | 0.053 |
|  | STG anterior | 0 | 0 | 0 | 0 | 0 | 0 |
|  | MTG anterior | 0 | 0 | 0 | 0 | 0 | 0 |
|  | STG posterior | 0 | 0 | 0 | 0 | 0 | 0 |
|  | MTG posterior | 0.121 | 0 | 0.102 | 0.157 | 0 | 0.121 |
|  | MTG temporal-occipital | 0.06 | 0 | 0 | 0 | 0.085 | 0.06 |
|  | Frontal pole | 0 | 0 | 0 | 0 | 0 | 0 |
|  | SFG | 0.01 | 0 | 0 | 0 | 0 | 0.01 |
|  | Central | 0 | 0 | 0 | 0 | 0 | 0 |
|  | Occipital pole | 0 | 0 | 0 | 0 | 0 | 0 |
|  | Perilesional 0-5 mm | 0.129 | 0.118 | 0 | 0.167 | 0 | 0.058 |
|  | Perilesional 5-10 mm | 0 | 0 | 0 | 0 | 0 | 0 |
|  | Perilesional 10-15 mm | 0 | 0 | 0 | -0.09 | 0 | 0 |
| **Lesion load** | IFG triangular | 0 | 0 | 0 | 0 | 0 | 0 |
|  | IFG opercularis | 0 | 0 | 0.184 | 0.121 | 0 | 0 |
|  | Supramarginal anterior | 0 | 0 | 0 | 0 | 0 | 0 |
|  | Supramarginal posterior | -0.217 | 0 | -0.297 | -0.139 | -0.203 | -0.217 |
|  | Angular gyrus | 0 | -0.047 | 0 | 0 | -0.01 | 0 |
|  | Temporal Pole | -0.009 | 0 | -0.156 | -0.055 | 0 | -0.009 |
|  | STG anterior | 0 | 0 | -0.125 | 0 | 0 | 0 |
|  | MTG anterior | 0 | 0 | -0.023 | 0 | 0 | 0 |
|  | STG posterior | 0 | 0 | 0 | -0.097 | 0 | 0 |
|  | MTG posterior | 0 | 0 | -0.033 | 0 | 0 | 0 |
|  | MTG temporal-occipital | -0.129 | 0 | -0.127 | -0.506 | -0.123 | -0.129 |
|  | Frontal Pole | -0.098 | 0 | -0.022 | 0 | 0 | -0.098 |
|  | SFG | 0 | 0 | 0 | 0 | 0 | 0 |
|  | Central | 0 | 0 | 0 | 0 | 0 | 0 |
|  | Occipital Pole | -0.082 | 0 | -0.254 | 0 | 0 | -0.082 |
| **Lambda** |  | 0.128 | 0.078 | 0.515 | 0.05 | 0.061 | 0.221 |
| **R<sup>2</sup>** |  | 0.567 | 0.738 | 0.167 | 0.806 | 0.722 | 0.538 |
| **RMSE** |  | 0.65 | 0.506 | 0.902 | 0.436 | 0.521 | 0.672 |

Regarding the lesion load covariate, we observed that greater lesion load was associated with lower language scores, particularly in perisylvian ROIs (hence, the negative coefficients). The only exception to this was the positive relationship between lesion load to the IFG opercularis and language outcomes. This pattern likely indicates that lesions specifically to that frontal area led to less severe language deficits, as critical posterior regions were spared in those instances (for a similar argument see Zhong et al.^46^).

## 4. DISCUSSION

In the current study, we first investigated how perfusion outside of lesioned areas was affected in chronic post-stroke aphasia in comparison to perfusion in age-matched controls. We then determined how perfusion in specific cortical and perilesional areas was related to language outcomes in aphasia. To investigate perfusion in aphasia we used anatomically defined ROIs from the Harvard-Oxford atlas that covered the frontal, parietal, and temporal areas of the perisylvian cortex in both hemispheres, along with several control regions not implicated previously in language processing. For the PWA group, we also analyzed perfusion levels in tissue at different distances from the lesion. We compared perfusion levels between the PWA and controls groups and investigated the relationship between perfusion levels and WAB subtest scores using both correlation and regularized regression analyses.

First, the current study demonstrated that cerebral perfusion in chronic stroke is greatly reduced even beyond the lesion site, with overall whole-brain perfusion significantly lower in the aphasia group compared to age-matched controls. The reduction in CBF is primarily noticeable in the lesioned hemisphere, with raw perfusion reduced in most regions in the left hemisphere and some regions in the contralesional hemisphere. Once ROI-specific perfusion values were normalized by the individual’s whole-brain perfusion, these adjusted perfusion levels in PWA still remained significantly reduced compared to controls in specific frontal and parietal areas outside of the lesion in the left lesioned hemisphere, while no statistically significant differences between PWA and controls were detected in homologous right hemisphere regions. This reduction in ipsilesional (left hemisphere) perfusion in areas distant from the lesion site has been observed in previous studies in both acute and chronic stroke using initially Positron Emission Tomography and more recently MRI perfusion-weighted imaging.^5,15,16,18–20,25,47,48^ However, we did not observe a pattern of hyperperfused right hemisphere regions, as found by Thompson and colleagues.^16^ Perilesional tissue closer to the lesion, as expected, showed lower perfusion compared to more distant perilesional areas, as has been documented in previous studies.^16–19^ Outside of the 5 mm tissue around the lesion, perfusion values returned to normal and were comparable to the rest of the brain. Furthermore, perfusion levels in the left hemisphere ROIs in PWA were lower compared to homologous right hemisphere regions, confirming previous findings.^16,25^ A similar trend of interhemispheric differences was observed for healthy controls, although to a lesser extent and the difference between perfusion in left and right hemispheres was only significant for four ROIs. Finally, asymmetry of perfusion between homologous left and right ROIs was more pronounced in the PWA group compared to the control group for frontal and parietal regions. This further supports the observation that greater decline in perfusion specifically in these ipsilateral areas is present in the aphasia group.

Second, we found that perfusion in the left temporal lobe (and most strongly in the posterior part of both superior and middle temporal gyri) and in the inferior parietal areas (supramarginal gyrus) was significantly related to different WAB subtest scores. This relationship was present even when direct lesion damage to these areas was accounted for, both in the partial correlation and the regularized regression analyses. This result indicates that spared blood flow in those areas is important for supporting language function. However, even if that tissue is preserved (as evidenced by an absence of a lesion on structural scans) it may display varying levels of functionality. Suboptimal CBF can affect function via impaired neurovascular coupling. CBF might be no longer matched to the metabolic requirements of the tissue and thus sub-ptimally perfused critical language areas might not be able to support residual processing. In contrast, perfusion in the frontal ROIs did not show such a relationship. This pattern is similar to a recent small study of perfusion in stroke patients, where perfusion levels – specifically in temporoparietal areas – were related to residual language abilities.^5^ Further, our findings are aligned with results seen by Thompson et al.^16^, although their correlations between perfusion and language scores did not survive correction for multiple comparisons. As expected, no relationship with language was observed for perfusion levels in control areas within the ipsilesional hemisphere.

In the perilesional analysis, only perfusion in the band of 0-5 mm was marginally related to language production abilities, particularly measures of fluency, but it did not survive the correction for multiple comparisons. In the regularized regression analysis only perilesional perfusion in the 0-5 mm band was again related to language production abilities, including fluency and naming ability, and overall aphasia severity. Thus, out of all the perilesional bands that we investigated only perfusion in tissue directly adjacent to the lesion was pertinent for language outcomes. More distant perilesional ROIs (5-10 mm and 10-15 mm) likely encompass different cytoarchitectonic areas that are unable to compensate properly for the lesioned tissue. Furthermore, aligned with most prior investigations, no relationship between perfusion in the contralesional (right hemisphere) ROIs and language abilities was observed.^16,25^

Overall, the results of the partial correlation analyses and the regularized regression were in concordance with each other. Areas identified in the partial correlation analyses were also shown to be predictive of language outcomes in the regularized regression. The uniqueness of the regularized regression is that it shows the joint contribution of multiple areas at the same time, highlighting which areas work together to support residual language abilities. All the language abilities, apart from repetition, were well predicted with the regularized regression models. It is possible that repetition, amongst all the language abilities tested in this study, is the most focal one and also relies more heavily on white matter integrity^49^, something that was not accounted for in the current study.

Collectively, these results argue that, as the language system reorganizes itself, the initial functional specialization of a region might be more pertinent than proximity to the lesion, with functionality of core temporoparietal language areas in the left hemisphere being most critical for determining residual language deficits. In other words, perfusion levels in language areas, rather than in perilesional tissue, determined the level of residual language abilities. Surprisingly no consistent strong associations were found between perilesional perfusion and post-stroke language abilities. This finding is likely due to the fact that participants were scanned in the chronic stage of their aphasia, where changes in the perilesional space have already taken place.^50,51^ It may also be the case that perilesional perfusion is more relevant in cases of motor recovery, as the motor function is more modular and spatially-restricted, i.e. more distant areas are not capable of taking on functions of the motor cortex.^52^

The current cross-sectional study was limited to exploring the role of perfusion in chronic stroke. Future investigations should explore changes in perfusion as individuals regain their language abilities in the first months post-stroke to better understand its causal and predictive role in stroke recovery. Additionally, while the current investigation took a significant step forward in terms of investigating the differential contributions of perfusion to language outcomes while accounting for structural cortical damage, the possible impact of structural disconnections on behavioral outcomes was not assessed. This will need to be comprehensively addressed in future multimodal neuroimaging studies. Further, more advanced perfusion imaging sequences, particularly those that use multiple post labeling delays, will allow more precise estimates of CBF while accounting for possible vascular delays.

### Conclusions

The current study was the largest exploration of perfusion in chronic post-stroke aphasia to date. Most importantly, we comprehensively investigated the relationship between perfusion in multiple regions simultaneously and language abilities while accounting for structural cortical damage, allowing us to uncover the unique contribution of residual perfusion to language outcomes. Overall, the results demonstrate that blood flow is reduced beyond the lesion site in chronic post-stroke aphasia and hypoperfused neural tissue in critical temporoparietal language areas is not fully able to support recovery leading to more pronounced residual language deficits. The findings underscore the critical and general role that left hemisphere posterior temporal regions play in various expressive and receptive language abilities.^49,53–55^ Overall, the study shows that slowed or reduced blood distribution can affect the functionality of regions beyond the lesion site and have a direct impact on behavioral outcomes.

The current study highlights the importance of exploring CBF measures in stroke. Perfusion MRI can be used alongside functional MRI in a complementary manner as it offers biologically meaningful quantitative measurement of tissue metabolism and function. Relative to task-based functional MRI, perfusion offers insights into functionality of all the brain regions simultaneously, not just those involved in the execution of a given task. Furthermore, a perfusion protocol can be easier to administer, and can be more readily standardized across sites and hence included in routine clinical assessments. Perfusion measures, while rarely implemented in multimodal neuroimaging studies (with a notable exception by Kristinsson et al.^56^) may offer valuable prognostic indicators of recovery potential and should be routinely included in future studies investigating the neural mechanisms of post-stroke recovery.

## ACKNOWLEDGEMENTS

We would like acknowledge Brian Curran for his help with collection of neuroimaging data. We also extend our thanks to Dr. Robert Knight for his contribution to the anatomical accuracy of lesion reconstructions. We would like to thank Dr. John Detre for supplying the stack-of-spirals pCASL sequence. As always, we are deeply grateful to all the participants with aphasia who make this research possible.

## REFERENCES

1. Markus HS. Cerebral perfusion and stroke. J Neurol Neurosurg Psychiatry. 2004;75(3):353–361. doi:10.1136/jnnp.2003.025825

2. Leenders KL, Perani D, Lammertsma AA, et al. Cerebral blood flow, blood volume and oxygen utilization: Normal values and effect of age. Brain. 1990;113(1):27–47. doi:10.1093/brain/113.1.27

3. Parkes LM, Rashid W, Chard DT, Tofts PS. Normal Cerebral Perfusion Measurements Using Arterial Spin Labeling: Reproducibility, Stability, and Age and Gender Effects. Magn Reson Med. 2004;51(4):736–743. doi:10.1002/mrm.20023

4. Joris PJ, Mensink RP, Adam TC, Liu TT. Cerebral blood flow measurements in adults: A review on the effects of dietary factors and exercise. Nutrients. 2018;10(5):1–15. doi:10.3390/nu10050530

5. Abbott NT, Baker CJ, Chen C, Liu TT, Love TE. Defining hypoperfusion in chronic aphasia: An individualized thresholding approach. Brain Sci. 2021;11(4):491. doi:10.3390/brainsci11040491

6. Newberg AB, Wang J, Rao H, et al. Concurrent CBF and CMRGlc changes during human brain activation by combined fMRI-PET scanning. Neuroimage. 2005;28(2):500–506. doi:10.1016/j.neuroimage.2005.06.040

7. Buxton RB, Uludağ K, Dubowitz DJ, Liu TT. Modeling the hemodynamic response to brain activation. Neuroimage. 2004;23(SUPPL. 1):220–233. doi:10.1016/j.neuroimage.2004.07.013

8. Astrup J, Symon L, Branston NM, Lassen NA. Cortical evoked potential and extracellular k+ and h+ at critical levels of brain ischemia. Stroke. 1977;8(1):51–57. doi:10.1161/01.STR.8.1.51

9. Sekhon LHS, Morgan MK, Spence I, Weber NC. Chronic cerebral hypoperfusion and impaired neuronal function in rats. Stroke. 1994;25(5):1022–1027. doi:10.1161/01.STR.25.5.1022

10. Powers WJ, Press GA, Grubb RL, Gado M, Raichle ME. The effect of hemodynamically significant carotid artery disease on the hemodynamic status of the cerebral circulation. Ann Intern Med. 1987;106(1):27–35. doi:10.7326/0003-4819-106-1-27

11. Feeney DM, Baron JC. Diaschisis. Stroke. 1986;17(5):817–830.

12. Girouard H, Iadecola C. Neurovascular coupling in the normal brain and in hypertension, stroke, and Alzheimer disease. J Appl Physiol. 2006;100(1):328–335. doi:10.1152/japplphysiol.00966.2005

13. Chalela JA, Alsop DC, Gonzalez-Atavales JB, Maldjian JA, Kasner SE, Detre JA. Magnetic resonance perfusion imaging in acute ischemic stroke using continuous arterial spin labeling. Stroke. 2000;31(3):680–687. doi:10.1161/01.STR.31.3.680

14. Demeestere J, Wouters A, Christensen S, Lemmens R, Lansberg MG. Review of perfusion imaging in acute ischemic stroke: From time to tissue. Stroke. 2020:1017–1024. doi:10.1161/STROKEAHA.119.028337

15. Hillis AE, Barker PB, Beauchamp NJ, Gordon B, Wityk RJ. MR perfusion imaging reveals regions of hypoperfusion associated with aphasia and neglect. Neurology. 2000;55(6):728–8. doi:doi: 10.1212/wnl.55.6.782.

16. Thompson CK, Walenski M, Chen Y, et al. Intrahemispheric perfusion in chronic stroke-induced aphasia. Neural Plast. 2017;(Article ID 2361619):1–15. doi:10.1155/2017/2361691

17. Boukrina O, Barrett AM, Graves WW. Cerebral perfusion of the left reading network predicts recovery of reading in subacute to chronic stroke. Hum Brain Mapp. 2019;(40):5301–5314. doi:10.1002/hbm.24773

18. Brumm KP, Perthen JE, Liu TT, Haist F, Ayalon L, Love T. An Arterial Spin Labeling investigation of cerebral blood flow deficits in chronic stroke survivors. Neuroimage. 2011;51(3):995–1005. doi:10.1016/j.neuroimage.2010.03.008.An

19. Richardson JD, Baker JM, Morgan PS, Rorden C, Bonilha L, Fridriksson J. Cerebral perfusion in chronic stroke: Implications for lesion-symptom mapping and functional MRI. Behav Neurol. 2011;24(2):117–122. doi:10.3233/BEN-2011-0283

20. Hillis AE. Magnetic resonance perfusion imaging in the study of language. Brain Lang. 2007;102(2):165–175. doi:10.1016/j.bandl.2006.04.016

21. Motta M, Ramadan A, Hillis AE, Gottesman RF, Leigh R. Diffusion – perfusion mismatch: An opportunity for improvement in cortical function. Front Neurol. 2015;5(280):1–8. doi:10.3389/fneur.2014.00280

22. Robson H, Specht K, Beaumont H, et al. Arterial spin labelling shows functional depression of non-lesion tissue in chronic Wernicke’s aphasia. Cortex. 2017;92:249–260. doi:10.1016/j.cortex.2016.11.002

23. Wiest R, Abela E, Missimer J, et al. Interhemispheric cerebral blood flow balance during recovery of motor hand function after ischemic stroke-A longitudinal MRI study using arterial spin labeling perfusion. PLoS One. 2014;9(9). doi:10.1371/journal.pone.0106327

24. Fridriksson J, Richardson JD, Fillmore P, Cai B. Left hemisphere plasticity and aphasia recovery. Neuroimage. 2012;60(2):854–863. doi:10.1016/j.neuroimage.2011.12.057

25. Mimura M, Kato M, Kato M, et al. Prospective and retrospective studies of recovery in aphasia. Changes in cerebral blood flow and language functions. Brain. 1998;121(11):2083–2094. doi:10.1093/brain/121.11.2083

26. Thompson CK, Ouden D Den, Bonakdarpour B, Garibaldi K, Parrish TB. Neural plasticity and treatment-induced recovery of sentence processing in agrammatism. Neuropsychologia. 2010;48(11):3211–3227. doi:10.1016/j.neuropsychologia.2010.06.036

27. Yarkoni T, Poldrack RA, Nichols TE, Van Essen DC, Wager TD. Large-scale automated synthesis of human functional neuroimaging data. Nat Methods. 2011;8(8):665–670. doi:10.1038/nmeth.1635

28. Oldfield RC. The assessment and analysis of handedness: The Edinburgh inventory. Neuropsychologia. 1971;9(1):97–113.

29. Kertesz A. Western Aphasia Battery. New York: Grune & Stratton; 1982.

30. Kertesz A. Western Aphasia Batterys - Revised. San Antonio, TX: PsychCorp; 2007.

31. Alsop DC, Detre JA, Golay X, et al. Recommended implementation of arterial spin-labeled Perfusion mri for clinical applications: A consensus of the ISMRM Perfusion Study group and the European consortium for ASL in dementia. Magn Reson Med. 2015;73(1):102–116. doi:10.1002/mrm.25197

32. Chang YV, Vidoretta M, Wang Z, Detre JA. 3D-Accelerated, Stack-of-spirals Acquisitions and Reconstruction of Arterial Spin Labeling MRI. Magn Reson Imaging. 2017;78(4):1405–1419. doi:10.1002/mrm.26549.3D-Accelerated

33. Jenkinson M, Beckmann CF, Behrens TEJ, Woolrich MW, Smith SM. FSL. Neuroimage. 2012;62(2):782–790. doi:10.1016/j.neuroimage.2011.09.015

34. Chappell MA, Groves AR, Whitcher B, Woolrich MW. Variational Bayesian inference for a nonlinear forward model. IEEE Trans Signal Process. 2009;57(1):223–236. doi:10.1109/TSP.2008.2005752

35. Rorden C, Brett M. Stereotaxic Display of Brain Lesions. Behav Neurol. 2000;12(4):191–200. doi:10.1155/2000/421719

36. Avants BB, Tustison NJ, Song G, Cook PA, Klein A, Gee JC. A reproducible evaluation of ANTs similarity metric performance in brain image registration. Neuroimage. 2011;54(3):2033–2044. doi:10.1016/j.neuroimage.2010.09.025

37. Fonov V, Evans A, McKinstry R, Almli C, Collins D. Unbiased nonlinear average age-appropriate brain templates from birth to adulthood. Neuroimage. 2009;47:S102. doi:10.1016/S1053-8119(09)70884-5

38. Desikan RS, Ségonne F, Fischl B, et al. An automated labeling system for subdividing the human cerebral cortex on MRI scans into gyral based regions of interest. Neuroimage. 2006;31(3):968–980. doi:10.1016/j.neuroimage.2006.01.021

39. R Core Team. R: A Language and Environment for Statistical Computing. 2020. https://www.r-project.org/.

40. Wickham H. Ggplot2: Elegant Graphics for Data Analysis. Springer-Verlag New York; 2016. https://ggplot2.tidyverse.org.

41. Kim S. ppcor: An R Package for a Fast Calculation to Semi-partial Correlation Coefficients. Commun Stat Appl Methods. 2015;22(6):665–674. doi:10.5351/csam.2015.22.6.665

42. Salvalaggio A, de Filippo De Grazia M, Zorzi M, de Schotten MT, Corbetta M. Post-stroke deficit prediction from lesion and indirect structural and functional disconnection. Brain. 2020;143(7):2173–2188. doi:10.1093/brain/awaa156

43. Hastie T, Tibshirani R, Friedman J. The Elements of Statistical Learning. 2nd ed. Springer Science+Business Media, LLC; 2009. https://hastie.su.domains/ElemStatLearn/printings/ESLII_print12_toc.pdf.

44. Tibshirani R, Bien J, Friedman J, et al. Strong rules for discarding predictors in lasso-type problems. J R Stat Soc. 2010;74(2):245–266.

45. Friedman J, Hastie T, Tibshirani R. Regularization paths for generalized linear models via coordinate descent. J Stat Softw. 2010;33(1):1–22. doi:10.18637/jss.v033.i01

46. Zhong AJ, Baldo J V., Dronkers NF, Ivanova M V. The unique role of the frontal aslant tract in speech and language processing. NeuroImage Clin. 2022;34(March):103020. doi:10.1016/j.nicl.2022.103020

47. Guadagno J V., Warburton EA, Jones PS, et al. The diffusion-weighted lesion in acute stroke: Heterogeneous patterns of flow/metabolism uncoupling as assessed by quantitative positron emission tomography. Cerebrovasc Dis. 2005;19(4):239–246. doi:10.1159/000084087

48. Sobesky J, Zaro Weber O, Lehnhardt FG, et al. Which time-to-peak threshold best identifies penumbral flow? A comparison of perfusion-weighted magnetic resonance imaging and positron emission tomography in acute ischemic stroke. Stroke. 2004;35(12):2843–2847. doi:10.1161/01.STR.0000147043.29399.f6

49. Ivanova M V., Zhong A, Turken A, Baldo J V., Dronkers NF. Functional contributions of the arcuate fasciculus to language processing. Front Hum Neurosci. 2021;15(June):1–15. doi:10.3389/fnhum.2021.672665

50. Lee RG, Donkelaar P Van. Mechanisms Underlying Functional Recovery Following Stroke. Can J Neurol Sci / J Can des Sci Neurol. 1995;22(4):257–263. doi:10.1017/S0317167100039445

51. Nudo RJ. Recovery after brain injury: mechanisms and principles. Front Hum Neurosci. 2013;7(December):1–14. doi:10.3389/fnhum.2013.00887

52. Nudo RJ. Recovery after damage to motor cortical areas. Curr Opin Neurobiol. 1999;9:740–747.

53. Wilson SM, Entrup JL, Schneck SM, et al. Recovery from aphasia in the first year after stroke. Brain. 2022.

54. Ivanova M V., Isaev DY, Dragoy O V., et al. Diffusion-tensor imaging of major white matter tracts and their role in language processing in aphasia. Cortex. 2016;85:165–181. doi:10.1016/j.cortex.2016.04.019

55. Turken AU, Dronkers NF. The Neural Architecture of the Language Comprehension Network: Converging Evidence from Lesion and Connectivity Analyses. Front Syst Neurosci. 2011;5(February):1–20. doi:10.3389/fnsys.2011.00001

56. Kristinsson S, Zhang W, Rorden C, et al. Machine learning-based multimodal prediction of language outcomes in chronic aphasia. Hum Brain Mapp. 2021;42(6):1682–1698. doi:10.1002/hbm.25321

