## Appendix for "Cerebral perfusion in post-stroke aphasia and its relation to residual language abilities"

Table A1. *Descriptive statistics (Mean (SD)) of mean raw perfusion for left and right hemispheres ROIs in the aphasia and the control groups along with results of between group comparison (aphasia vs. control group). Significant tests (p < .0033) are bolded.*

| ROI | Aphasia | | Controls | | Left hemisphere comparisons | | | Right hemisphere comparisons | | |
|---|---|---|---|---|---|---|---|---|---|---|
| | LH | RH | LH | RH | Test | statistic | p-value | Test | statistic | p-value |
| IFG triangular | 25.77 (13.89) | 31.43 (11.44) | 36.36 (8.71) | 39.34 (9.96) | **Welch-test** | **-3.86** | **0.0002597** | T-test | -2.88 | 0.0053782 |
| IFG opercularis | 25.31 (13.13) | 35.29 (11.43) | 40.3 (9.66) | 42.68 (9.41) | **T-test** | **-4.96** | **0.0000054** | T-test | -2.74 | 0.0079305 |
| Supramarginal gyrus anterior | 19.92 (9.72) | 26 (8.96) | 29.73 (7.25) | 29.73 (5.78) | **T-test** | **-4.38** | **0.0000433** | Wilcox-test | 381.00 | 0.0472330 |
| Supramarginal gyrus posterior | 22.32 (9.73) | 29.42 (8.88) | 33.93 (6.64) | 37.34 (7.2) | **Welch-test** | **-5.83** | **0.0000002** | **T-test** | **-3.79** | **0.0003307** |
| Angular gyrus | 23.55 (10.14) | 29.67 (9.44) | 36.48 (6.75) | 38.31 (7.84) | **Wilcox-test** | **165.00** | **0.0000022** | **T-test** | **-3.87** | **0.0002553** |
| Temporal Pole | 22.93 (10.8) | 28.71 (11.7) | 32.58 (9.08) | 34.71 (10.23) | **T-test** | **-3.76** | **0.0003671** | T-test | -2.13 | 0.0367816 |
| STG anterior | 28.74 (12.92) | 36.23 (12.39) | 38.45 (8.07) | 39.83 (9.97) | **Welch-test** | **-3.76** | **0.0003723** | Wilcox-test | 446.00 | 0.2470875 |
| MTG anterior | 22.98 (13.09) | 24.62 (12.08) | 30.58 (11.29) | 31.45 (10.03) | Wilcox-test | 334.00 | 0.0135243 | T-test | -2.39 | 0.0198897 |
| STG posterior | 28.42 (10.51) | 37.5 (10.74) | 40.5 (9.63) | 48.79 (9.53) | **T-test** | **-4.67** | **0.0000158** | **Wilcox-test** | **228.00** | **0.0000849** |
| MTG posterior | 26.06 (12.34) | 32.37 (11.35) | 33.97 (8.96) | 38.66 (9.28) | T-test | -2.80 | 0.0066482 | Wilcox-test | 333.00 | 0.0094666 |
| MTG temporal-occipital | 28.13 (11.18) | 35.63 (11.55) | 37.15 (7.55) | 41.2 (8.69) | **Welch-test** | **-3.96** | **0.0001904** | Wilcox-test | 344.00 | 0.0140956 |
| Frontal Pole | 18.5 (8.81) | 22.33 (8.07) | 28.28 (7.98) | 30.45 (7.26) | **T-test** | **-4.57** | **0.0000221** | **Wilcox-test** | **231.00** | **0.0000994** |
| SFG | 22.49 (9.45) | 24.22 (8.27) | 27.82 (8.57) | 27.38 (9.43) | T-test | -2.32 | 0.0235239 | T-test | -1.44 | 0.1539328 |
| Central | 22.51 (10) | 27.54 (8.72) | 31.23 (6.12) | 33.24 (6.13) | **Welch-test** | **-4.46** | **0.0000326** | T-test | -2.88 | 0.0053929 |
| Occipital Pole | 22.15 (9.75) | 26.34 (12) | 30.3 (7.94) | 33.15 (9.04) | **Wilcox-test** | **245.00** | **0.0002040** | **Wilcox-test** | **279.00** | **0.0010323** |

Table A2. *Descriptive statistics (Mean (SD)) of mean adjusted perfusion for left and right hemispheres ROIs in the aphasia and the control groups along with results of between group comparison (aphasia vs. control group). Significant tests (p < .0033) are bolded.*

| ROI | Aphasia | | Controls | | Left hemisphere comparisons | | | Right hemisphere comparisons | | |
|---|---|---|---|---|---|---|---|---|---|---|
| | LH | RH | LH | RH | Test | statistic | p-value | Test | statistic | p-value |
| IFG triangular | 0.94 (0.35) | 1.21 (0.31) | 1.12 (0.19) | 1.21 (0.19) | Wilcox-test | 364.00 | 0.0277763 | Wilcox-test | 541.00 | 0.9695619 |
| IFG opercularis | 0.92 (0.33) | 1.37 (0.31) | 1.25 (0.21) | 1.32 (0.17) | **Welch-test** | **-4.99** | **0.0000049** | Welch-test | 0.91 | 0.3645874 |
| Supramarginal gyrus anterior | 0.75 (0.27) | 1.01 (0.27) | 0.92 (0.14) | 0.92 (0.11) | Wilcox-test | 315.00 | 0.0047476 | Welch-test | 1.93 | 0.0579147 |
| Supramarginal gyrus posterior | 0.84 (0.22) | 1.14 (0.22) | 1.05 (0.13) | 1.16 (0.11) | **Wilcox-test** | **208.00** | **0.0000286** | Welch-test | -0.28 | 0.7806069 |
| Angular gyrus | 0.89 (0.22) | 1.15 (0.24) | 1.13 (0.13) | 1.18 (0.1) | **Wilcox-test** | **149.00** | **0.0000008** | Welch-test | -0.83 | 0.4098371 |
| Temporal Pole | 0.88 (0.38) | 1.1 (0.31) | 1 (0.19) | 1.06 (0.22) | Wilcox-test | 386.00 | 0.0547811 | T-test | 0.50 | 0.6156078 |
| STG anterior | 1.13 (0.47) | 1.41 (0.4) | 1.2 (0.18) | 1.23 (0.24) | Welch-test | -0.75 | 0.4558823 | Welch-test | 2.29 | 0.0254819 |
| MTG anterior | 0.88 (0.41) | 0.92 (0.37) | 0.93 (0.24) | 0.96 (0.21) | Welch-test | -0.70 | 0.4848236 | Welch-test | -0.59 | 0.5594751 |
| STG posterior | 1.15 (0.42) | 1.47 (0.3) | 1.25 (0.17) | 1.52 (0.23) | Welch-test | -1.40 | 0.1670569 | T-test | -0.70 | 0.4838612 |
| MTG posterior | 1.01 (0.39) | 1.25 (0.27) | 1.05 (0.17) | 1.19 (0.16) | Wilcox-test | 485.00 | 0.5083549 | Welch-test | 1.06 | 0.2913353 |
| MTG temporal-occipital | 1.1 (0.36) | 1.38 (0.28) | 1.15 (0.13) | 1.27 (0.16) | Welch-test | -0.80 | 0.4247602 | Welch-test | 2.03 | 0.0469028 |
| Frontal Pole | 0.68 (0.17) | 0.85 (0.15) | 0.87 (0.15) | 0.94 (0.11) | **T-test** | **-4.58** | **0.0000213** | T-test | -2.46 | 0.0165881 |
| SFG | 0.85 (0.23) | 0.94 (0.21) | 0.85 (0.16) | 0.83 (0.19) | Welch-test | 0.00 | 0.9987794 | T-test | 2.06 | 0.0436407 |
| Central | 0.84 (0.22) | 1.07 (0.22) | 0.97 (0.12) | 1.03 (0.1) | **Welch-test** | **-3.11** | **0.0027797** | Welch-test | 0.94 | 0.3510641 |
| Occipital Pole | 0.83 (0.2) | 0.99 (0.22) | 0.93 (0.16) | 1.02 (0.16) | Wilcox-test | 347.00 | 0.0156637 | T-test | -0.60 | 0.5509859 |

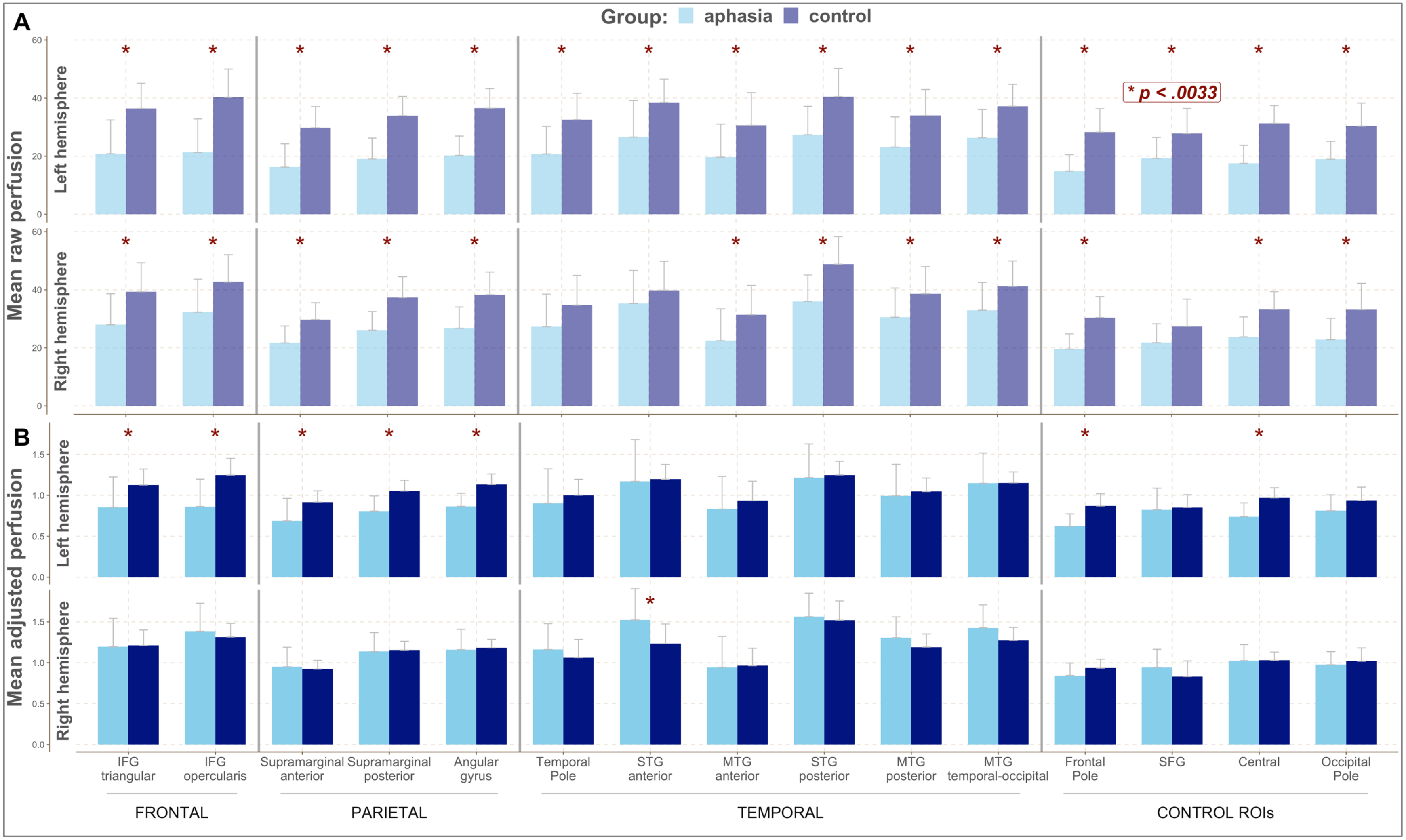


**Figure A1.** Mean perfusion values for the aphasia and the control groups from the VA cohort across left and right hemisphere ROIs. Red asterisks mark significant differences between groups for a given ROI. Panel A – Mean raw perfusion. Panel B – Mean adjusted perfusion. IFG – inferior frontal gyrus, STG – superior temporal gyrus, MTG – middle temporal gyrus, SFG – superior frontal gyrus.
